# Low-intensity focused ultrasound to the insula and dorsal anterior cingulate has site-specific and pressure dependent effects on pain during measures of central sensitization

**DOI:** 10.1101/2024.01.10.575098

**Authors:** Alexander In, Andrew Strohman, Brighton Payne, Wynn Legon

## Abstract

**Background:** The insula and dorsal anterior cingulate cortex (dACC) are core brain regions involved in pain processing and central sensitization, a shared mechanism across various chronic pain conditions. Methods to modulate these regions may serve to reduce central sensitization, though it is unclear which target may be most efficacious for different measures of central sensitization.

**Objective/Hypothesis:** Investigate the effect of low-intensity focused ultrasound (LIFU) pressure to the anterior insula (AI), posterior insula (PI) or dACC on conditioned pain modulation (CPM) and temporal summation of pain (TSP).

**Methods:** N = 16 volunteers underwent TSP and CPM pain tasks pre/post a 10 minute LIFU intervention to either the AI, PI, dACC or Sham stimulation. Pain ratings were collected pre/post LIFU.

**Results:** LIFU to the PI significantly attenuated pain ratings in both TSP and the CPM protocols. LIFU to the dACC only affected TSP pain ratings. LIFU to AI had no effect on either TSP or CPM pain ratings. LIFU pressure modulated group means but did not affect overall group differences.

**Conclusions:** LIFU to the PI and dACC differentially affected central sensitization. This may, in part, be due to dosing (pressure) of LIFU. Inhibition of the PI with LIFU may be a future potential therapy in chronic pain populations demonstrating central sensitization. The minimal effective dose of LIFU for efficacious neuromodulation will help to translate LIFU for therapeutic options.

## INTRODUCTION

Low-intensity focused ultrasound (LIFU) is a non-invasive neuromodulatory approach that uses mechanical energy to reversibly modulate neuronal activity with high spatial resolution and adjustable depth of focus [1,2]. LIFU has been extensively studied in small [3–5] and large animals [6–9] including humans [10–13]. It has been demonstrated to produce safe [14,15] and reversible inhibition and excitation [8,16] in different parts of the cortex [12,13,17] and sub-cortical structures [1,18] including emerging clinical applications [19–23]. An additional promising clinical application for LIFU is pain as demonstrated in both primates [8,24] and humans [25,26]. Chronic pain patients routinely demonstrate increased activity of both the insula and dorsal anterior cingulate cortex (dACC) in response to painful stimuli as compared to healthy control participants [27–29]. In addition, many chronic pain patients demonstrate central sensitization that has also been linked to both the insula and dACC [30].

Central sensitization (CS) is a pathophysiologic process which represents abnormal responsiveness of neurons in the nociceptive pathway to their normal or subthreshold input [30,31]. CS is characterized by the amplification of pain by the central nervous system and is believed to be an underlying mechanism of various chronic pain conditions including fibromyalgia, complex regional pain syndrome, and chronic low back pain [32,33]. These conditions show dysfunction in pain processing as indexed by laboratory measures of central sensitization, which include temporal summation of pain (TSP) and conditioned pain modulation (CPM) [34–38]. TSP is the perception of increased pain evoked by repetitive noxious stimuli intended to index bottom-up pain processing [39]. CPM tests the response to a painful test stimulus either during or immediately after a painful conditioning stimulus [40], thought to measure pain inhibitory pathways and index top-down pain processing [41,42]. Several studies employing TSP and CPM have demonstrated increased activity of critical brain regions involved in pain processing including the insula [30,43–46] and dACC [30,44,46,47].

The insula is part of the cerebral cortex, deep to the lateral sulcus, covered by the frontal operculum [48]. It is comprised of both an anterior (AI) and posterior (PI) lobule differing in both structural and functional connectivity [49–52]. The dACC is located dorsal to the genu of the corpus callosum, stretching rostrally to the frontopolar cortex and caudally to its border with the posterior midcingulate cortex [53–55]. The PI and dACC receive the majority of direct di-synaptic spinothalamic projections through the ventral medial and medial dorsal thalamic nuclei, respectively, and thus thought to be involved in bottom-up pain processing [56–61]. The AI has strong reciprocal connections with both PI and dACC and is believed to integrate these signals with emotional and cognitive aspects of pain to form the overall pain percept [60,62]. Together, the insula and dACC look to be core nodes of cortical pain processing [56,63–65].

It is the purpose of this study to investigate how LIFU to AI, PI, or dACC affect pain ratings during TSP or CPM tasks. We hypothesize LIFU will reduce pain ratings (AI, PI, and dACC) as compared to Sham stimulation, and more specifically that LIFU to PI and dACC will be specific to TSP (bottom-up pain processing) and that LIFU to AI will specifically affect CPM (top-down pain processing) due to their respective structural and functional connectivity and purported role in pain processing [58,60,62]. Because dACC directly receives nociceptive input but is also implicated in top-down cognitive and motivation aspects of pain [54], we expect LIFU to dACC to affect both TSP and CPM pain ratings.

## MATERIALS & METHODS

### Participants

The Institutional Review Board at Virginia Tech approved all experimental procedures (IRB #21-796). N=16 healthy participants (25.7 years ± 3.4 years; range (20-34); M/F 5/11), who met all inclusion/exclusion criteria provided written informed consent to all aspects of the study. Inclusion criteria were males and females ages 18-65 while exclusion criteria included contraindications to imaging (magnetic resonance imaging (MRI) and computed tomography (CT)), a history of neurologic disorder or head injury resulting in loss of consciousness for >10 minutes, drug dependence, any active medical disorder or current treatment with potential central nervous system effects, and a history of chronic pain.

### Overall Study Design and Timeline

All participants completed a total of five visits in this sham-controlled cross-over pre/post design study. Following informed consent, the first visit comprised both a structural brain MRI and CT used for LIFU targeting and acoustic modelling (see below). The remainder of the visits were formal testing sessions of LIFU to the AI, PI, dACC, or Sham, randomized within and between subjects. A minimum of two days was required between visits to mitigate any potential carry-over effects as evidence suggests LIFU effects can last for minutes up to several hours [66]. At each visit, participants completed a review of symptoms (ROS) questionnaire, state anxiety inventory (STAI-S), and a daily activities questionnaire that queried sleep, physical activity, caffeine, and any recreational drug use or prescription medication on the day of testing. The general outline of the testing session was as follows: five minutes of resting baseline followed by counterbalanced TSP and CPM protocols (see below for details). Each task took roughly 10 minutes and were separated by 5-minute rest periods. The LIFU or Sham intervention was then administered for 10 minutes and tasks were repeated post-intervention. Task orders pre and post intervention were counterbalanced within and between subjects. Thirty minutes after the end of LIFU stimulation, the ROS and auditory masking questionnaires (AMQ) were administered. Each visit lasted roughly 2.5 hours. Continuous electroencephalography (EEG), electrocardiography (ECG) and electrodermal activity (EDR) was recorded throughout the entire testing session (detail below). **See Figure 1** for an example of data collection and timing of tasks during a single visit.

**Figure 1.**
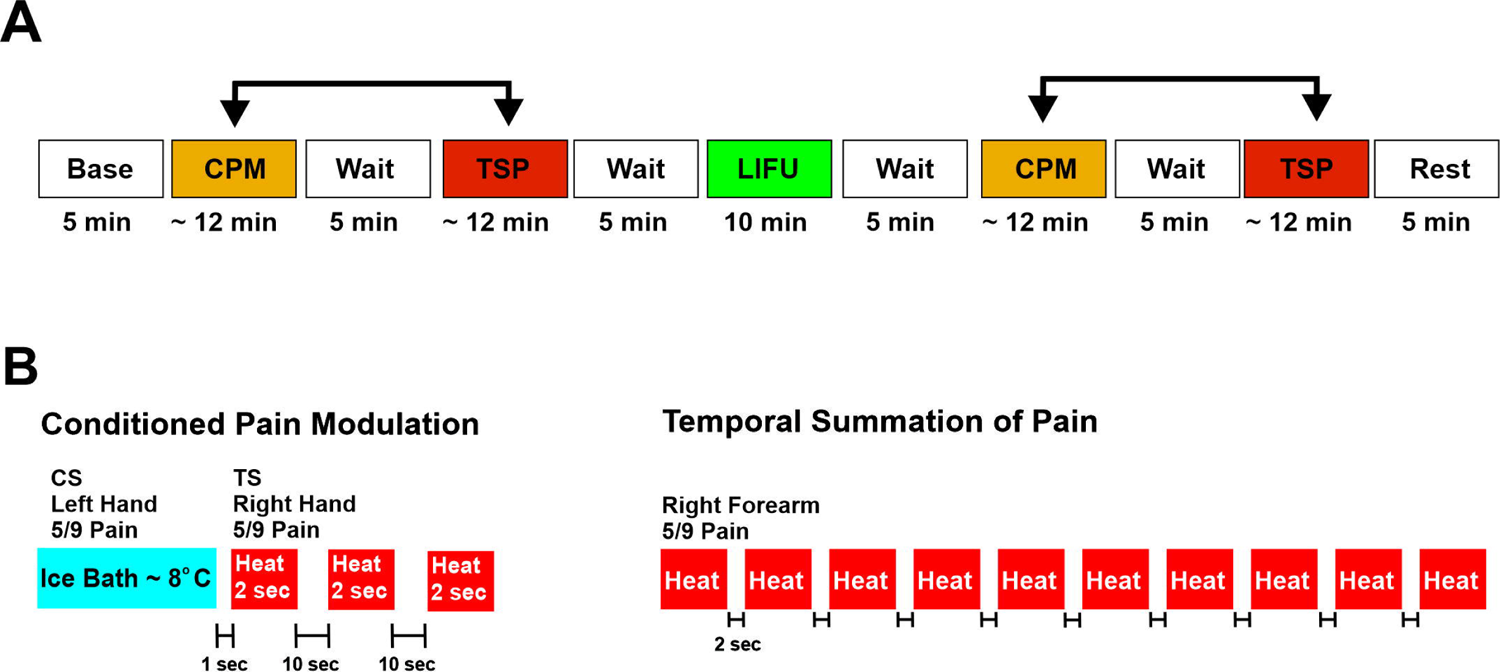
Overview of study collection and pain tasks. **A.** Pictorial representation of the timing of conditioned pain modulation (CPM) and temporal summation of pain (TSP) tasks relative to the low-intensity focused ultrasound (LIFU) intervention. Arrows represent that CPM and TSP order was counterbalanced within subject across sessions. **B.** (Left) Pictorial representation of CPM task. Participants were required to submerge their left hand above the wrist into an ice bath (the condition stimulus (CS)) with an average temperature of 8°C until they reached a subjective pain report of 5/9 at which point they removed their hand. The test stimulus (TS) was a 2 second heat stimulus (5/9 heat pain) delivered to the dorsum of the right hand. Three test stimuli were delivered immediately after removal of the CS and again at ten second intervals. Participants were required to verbally rate their pain from the TS which was used as the primary outcome measure pre/post LIFU. (Right) Pictorial representation of the TSP task. Ten heat stimuli of 1.4 seconds (previously titrated to a 5/9 pain) were delivered to the right forearm every two seconds. Participants were required to verbally rate each stimulus on a 0-9 pain scale.

### MRI and CT Acquisition

MRI data were acquired on a Siemens 3T Prisma scanner (Siemens Medical Solutions, Erlangen, Germany) at the Fralin Biomedical Research Institute’s Human Neuroimaging Laboratory. Anatomical scans were acquired using a T1-weighted MPRAGE sequence with a TR = 1400 ms, TI = 600 ms, TE = 2.66 ms, flip angle = 12 (degrees), voxel size = 0.5×0.5×1.0 mm, FoV read = 245 mm, FoV phase of 87.5%, 192 slices, ascending acquisition. CT scans were collected with a Kernel = Hr60 in the bone window, FoV = 250 mm, kilovolts (kV) = 120, rotation time = 1 second, delay = 2 seconds, pitch = 0.55, caudocranial image acquisition order, 1.0 mm image increments for a total of 121 images and scan time of 13.14 seconds.

### Pain Thresholding

Contact heat stimuli were delivered using a contact 3×3.2×2.4 mm peltier thermode (T03, QST.lab, Strasbourg, FR) and a thermal cutaneous stimulator (TCS, QST.lab, Strasbourg, FR). This device is capable of delivering rapid cooling or heating up to 300°C/s. This device has been validated in previous reports in both healthy volunteers [67–71] and clinical pain patients [72,73]. At each formal testing visit (sessions 2-5) participants were familiarized with the thermal probe prior to heat pain thresholding. Three methods of thresholding were conducted.

### Warm Thresholding

The thermode was placed 10 cm distally from the medial humeral epicondyle on a line from the medial epicondyle to the styloid process of the ulna over the muscle bulk of the wrist flexors as described in Mailloux et al. (2021) [74]. Starting at a baseline temperature of 30°C, the device increased temperature at a rate of 0.3°C/s, and the participant was required to press a button when they first felt a warm sensation. The button press also served to rapidly return the thermode to baseline temperature. This procedure was repeated 5 times with 2 seconds between each trial, and the average of the five stimuli was recorded as their warm threshold.

### Pain Onset Thresholding

Pain thresholding was performed on two body sites – the right hand dorsum and forearm. For the hand dorsum, the thermode was place over the capitate bone. For the forearm, the thermode was place as describe above in warm thresholding. For each site, the thermode baseline temperature was 30°C, and increased at a rate of 1°C/s. Participants were required to press a button when the sensation first became painful. The button press also served to rapidly return the thermode to baseline temperature. This procedure was repeated 5 times for each body site with a 30 second inter-stimulus interval. The order of body site testing was counterbalanced within and between subjects across sessions. The average of the five stimuli for each body site was recorded as their pain onset threshold.

### 5/9 Pain Thresholding

The thermode was placed on the participant’s hand and forearm as above. Starting at a baseline temperature of 30°C, the device increased temperature at a rate of 1°C/s. The participant was required to press a button when they first felt a 5/9 pain rating using a numerical pain rating scale (NPRS) from 0-9 with anchors 0 = no pain and 9 = unbearable pain. The NPRS had additional markings such that 1-3 = mild pain; 4-6 = moderate pain; 7-9 = severe pain. 5/9 thresholding was performed on both the arm and hand. This procedure was repeated five times on each body location with 30 seconds between each stimulus, and the average of the five stimuli was recorded as the participants’ mean 5/9 pain threshold for each body site.

### Temporal Summation of Pain (TSP)

TSP procedure was adapted from Staud et al. (2014) [37]. Prior to formal testing, the thermode was placed on the participant’s right forearm (as above) and started at 40°C then slowly cooled down (−1°C/sec) until reaching skin temperature. Once it reached skin temperature, 10 stimuli based on the participant’s 5/9 average threshold (for the forearm site) were administered. Each stimulus had a duration of 1.8 seconds that included a ramping up time at a rate of 70°C/s and a ramp down rate of 300°C/s. Because each participants’ 5/9 temperature was different, plateau time was slightly different but averaged ∼ 1.4 seconds. Each stimulus was delivered at a fixed inter-stimulus interval (ISI) of 2 seconds, returning to 40°C between each stimulus. Participants were required to verbally rate the intensity of each stimulus using the NPRS. This procedure was completed a total of three times, separated by 2 minutes. Total task time was roughly 10 minutes. This procedure was performed both pre and post intervention. **Figure 1B**. **Conditioned Pain Modulation (CPM)**

Conditioned pain modulation was performed according to recommendations of Yarnitsky et al. (2015) [40]. The conditioning stimulus (CS) was an ice bath with a temperature of 8°C. The test stimulus (TS) was the heat stimulus based on individual subject (and individual session) 5/9 average heat pain at the dorsum of the hand. The CS was delivered to the left hand and the TS to the right hand. Participants submerged their left hand up to the wrist in the ice water and were required to keep it in the ice bath until their pain reached a 5/9 using the NPRS. There was no minimum time requirement. Immediately upon removal of the left hand from the ice bath, three TS were delivered to the dorsum of the right hand at an ISI of 10 seconds. We followed the recommendation of Yarnitsky et al. (2015) [40] to perform the TS after the CS. The TS had a duration of 2 seconds including a ramp up time at a rate of 100°C/s and a ramp down rate of 300°C/s. Baseline temperature was set at 32°C. Participants were required to verbally rate the pain intensity of the TS using the NPRS. CPM was conducted three times with an ISI of 2 minutes. Total task time was roughly 12 minutes. This procedure was performed both pre and post intervention. **Figure 1B**.

### LIFU Transducers and Waveform

Two different transducers were used based upon individual participant head morphology and depth of target (Individual participant depth of targets is provided in **Supplementary Table 2**). For insula targets, we used a single-element 500 kHz transducer (Sonic Concepts H-281) with an active diameter of 45.0 mm and 38.0 mm focal length from the exit plane. The transducer also had a solid water coupling over the radiating surface to the exit plane. To target the dACC, we used a single-element 500 kHz transducer (Sonic Concepts H-104) with an active diameter of 64.0 mm and 52.0 mm focal length from the exit plane. See **Figure 2A** for free water empirical lateral and axial pressure plots.

**Figure 2.**
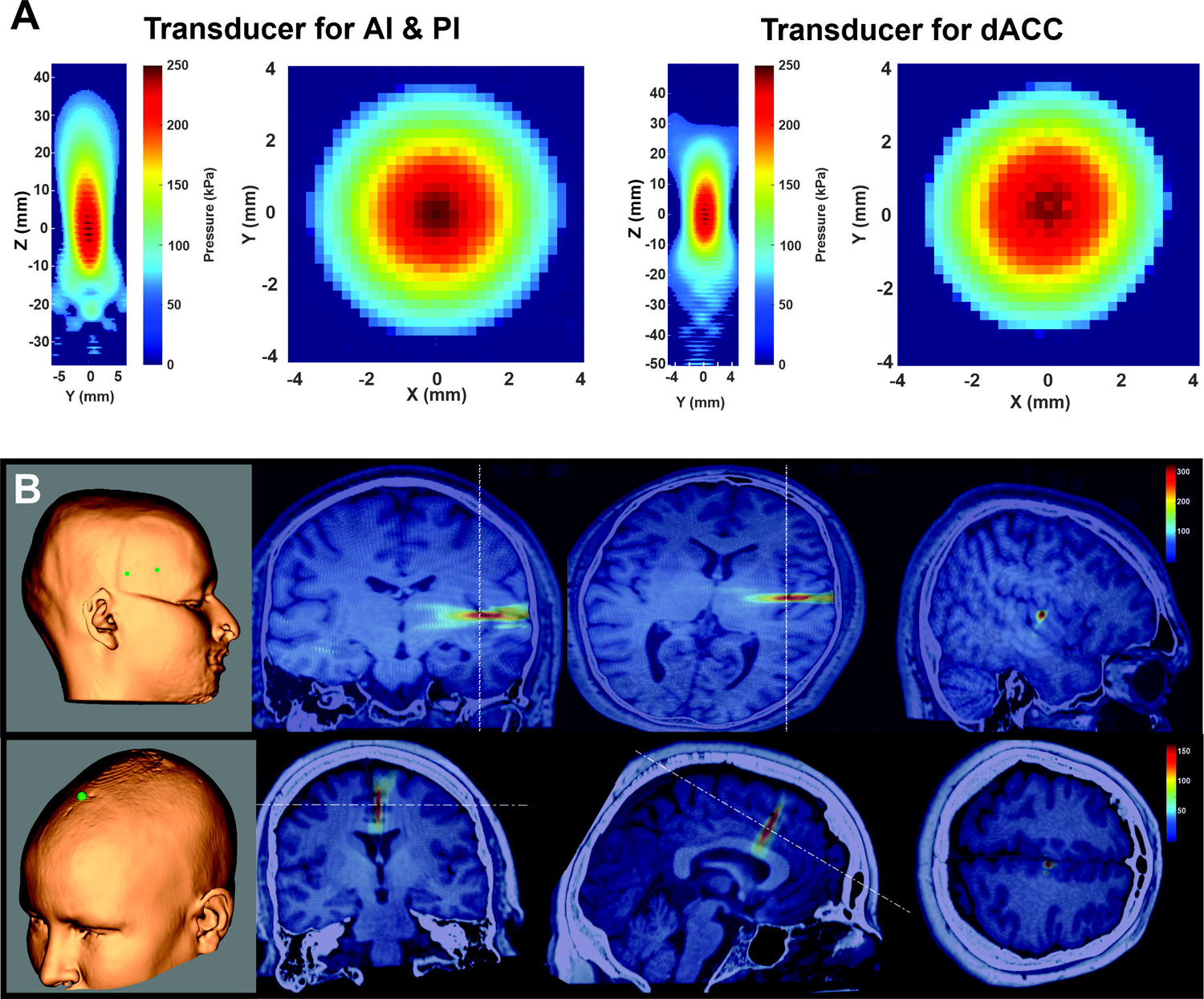
Ultrasound maps and models. **A.** Pseudocolor YZ and XY empirical measurements from an acoustic test tank of the pressure map for the transducer used for targeting anterior (AI) and posterior (PI) insula (left) and for the dorsal anterior cingulate cortex (dACC) (Right). kPa = kiloPascals. **B.** Scalp targets from neuronavigation system showing positioning of the center of the transducer for insular targets (top left) and dACC (bottom left). (Right) Acoustic models using individual magnetic resonance and computed tomography scans from representative subjects showing 500 kHz LIFU targeting the posterior insula (top) and dACC (bottom). Colorscale is kPa.

Focused ultrasound waveforms were generated using a dual channel function generator (BK 4078B Precision Instruments). Channel 1 was a 5Vp-p square wave burst of 1 kHz (N=1000) with a duty cycle of 36% used to gate channel 2 that was a 500 kHz sine wave. Channel 2 output was sent through a 100-W linear RF amplifier (E&I 2100L; Electronics & Innovation) before being sent to the transducers. LIFU was applied for 100 stimulations (burst duration = 1 sec) separated by a fixed ISI of 5 seconds for total application time of 10 minutes. For both AI and PI, the applied external pressure was 380 kPa with a spatial peak-pulse average intensity (I_sppa_) of 4.2 W/cm^2^, a spatial peak temporal average (I_spta_) of 1.5 W/cm^2^, and a mechanical index (MI) of 0.2. For dACC, the applied external pressure was 400 kPa with an I_sppa_ of 4.5 W/cm^2^, an I_spta_ of 1.62 W/cm^2^, and an MI of 0.23.

### LIFU Targeting and Application

The transducer was coupled to the head using conventional ultrasound gel and our custom mineral oil/polymer coupling pucks [75]. These pucks have negligible attenuation at 500 kHz [75] and can be made with varying stand-off heights that allow for precise axial (depth) targeting based on individual insular and dACC target depths. Each participant’s left AI and PI target was identified with the aid of an insular atlas [46] and depth was measured from the scalp. An appropriate coupling puck was made so that the focal spot of the transducer (38 mm) was exactly overlaid on the insular target. For dACC, MNI coordinate [0,18, 30] was used based upon Neurosynth reverse inference software for the term ‘pain’ and prior literature on the fMRI signature of heat pain [63,76]. Placement of the transducer on the scalp for each target site was aided using a neuronavigation system (BrainSight, Rogue Research, Montreal, QUE, CAN) with built in MNI-to-native space transformations for the dACC target. LIFU or Sham was only delivered if placement error on the scalp was < 3 mm. See **Figure 2B** for example of scalp targets for each of anterior and posterior insula and dACC.

### Auditory Masking

It is known that single-element transducers can produce an audible auditory artifact that can potentially confound the data [77,78] but has also been demonstrated to be effectively masked using appropriate masking stimuli [77,79,80]. Therefore, we utilized continuous acoustic masking to remove any audible sounds throughout the entire LIFU or sham application. Participants were given headphones that were connected to a Kindle tablet that had a multi-tone white noise generator app. There were a variety of sound options that could be mixed and layered on top of each other to create a multitoned sound that has been established to effectively mask a 1 kHz pulsed ultrasound application [79]. The volume was set to a level where participants could not hear normal conversations with the intensities ranging from 70 to 75 dB.

### Questionnaires

#### Review of Symptoms

A review of symptoms questionnaire[14] asks about the presence of various symptoms and their severity (absent / mild / moderate / severe) scored on a scale of 0-3. This questionnaire was collected at the beginning of each visit (sessions 2 – 5) and 30 minutes after the LIFU or sham application in order to indicate any changes of symptoms from the intervention.

#### Auditory Masking

An auditory masking questionnaire [75,81] was administered at the end of each visit (sessions 2 – 5) with the following questions: “I could hear the LIFU stimulation”, “I could feel the LIFU stimulation”, and “I believe I experienced LIFU stimulation.” Participants were asked to rate each question on a 7-point Likert scale: strongly disagree / disagree / somewhat disagree / neutral / somewhat agree / agree / strongly agree.

### Data Analysis

#### Acoustic Modelling

Computational models were developed using individual subject MR and CT images to evaluate the wave propagation of LIFU across the skull and the resultant intracranial acoustic pressure maps. Simulations were performed using the k-Wave MATLAB toolbox[82], which uses a pseudospectral time domain method to solve discretized wave equations on a spatial grid. CT images were used to construct the acoustic model of the skull, while MR images were used to target LIFU at either the AI, PI, or dACC target, based on individual brain anatomy. Details of the modeling parameters can be found in Legon et al. (2018)[1]. CT and MR images were first co-registered and then up-sampled for acoustic simulations and the acoustic parameters for simulation were calculated from the CT images. The skull was extracted manually using a threshold intensity value and the intracranial space was assumed to be homogenous as ultrasound reflections between soft tissues are small [83]. Acoustic parameters were calculated from CT data assuming a linear relationship between skull porosity and the acoustic parameters[84,85]. The computational model of the ultrasound transducer used in simulations was constructed to recreate empirical acoustic pressure maps of focused ultrasound transmitted in the acoustic test tank similar to previous work[1,10,81,86]. See **Figure 2B** for an acoustic model from one representative participant demonstrating targeting of posterior insula and dACC.

#### Temporal Summation of Pain

The pain ratings for each of the 10 stimulations were averaged across participants for both pre and post LIFU (or sham) sessions. The difference (post-pre) in pain ratings were calculated, providing 10 difference values for all conditions for each participant. Pain ratings differences (post-pre) were then averaged across all participants for each condition. A two-way repeated measures analysis of variance was conducted on the difference scores with main effects LIFU (AI, PI, dACC, Sham) and TIME (0 – 10 seconds) and LIFU*TIME interaction. Interactions and main effects were explored using suitable post-hoc analyses. Significance was set at p < 0.05.

### Conditioned Pain Modulation

The pain ratings for each of the three stimulations were averaged across participants for both pre and post LIFU or sham sessions. The pain rating differences were calculated between pre and post sessions (post-pre) for all conditions across all participants. A two-way repeated measures analysis of variance (ANOVA) was conducted with main effects LIFU (AI, PI, dACC, Sham) and TIME (0,10,20) and the LIFU*TIME interaction. Interaction and main effects were explored using suitable post-hoc testing with significance set at p < 0.05.

## RESULTS

### LIFU Beam Characteristics

#### Insula

For the H-281 transducer, the free water beam profile revealed a full-width half maximum (FWHM) lateral resolution of ±1.7 mm at Z maximum and FWHM axial resolution of ±7.5 mm with a focal depth of 38 mm from the exit plane of the transducer **Figure 2A**.

#### dACC

For the H-104 transducer, the free water beam profile revealed a FWHM lateral resolution of ±2.0 mm at Z maximum and FWHM axial resolution of ±10 mm with a focal depth of 52 mm from the exit plane of the transducer **Figure 2A**.

### Brain Targeting

The average depth ± SD from the scalp to the AI, PI, and dACC for males was: 36.1 ± 3.8, 41.2 ± 3.4, and 50.3 ± 4.1 mm. The average depth ± SD to the AI, PI, and dACC for females was: 34.1 ± 1.5, 38.9 ± 1.7, and 45.5 ± 2.1 mm. The mean for all participants was: 34.1 ± 1.5 mm, 39.0 ± 1.7 mm, and 45.5 ± 2.1 mm. This was in good agreement with the FWHM of each of the transducers used to target these regions. Individual participant depth of target is in **Table S1**. Acoustics models showing targeting of PI and dACC are shown in **Figure 2B**.

### Estimated intracranial pressure

The mean ± SD estimated intracranial pressure in the head for AI, PI, and dACC was: 180.1 ± 84.6 kPa, 262.1 ± 93.6 kPa, and 119.8 ± 55.7. The one-way repeated measures ANOVA revealed a main effect: F(2,30) = 17.81, p = 7.96e-06. Post-hoc Tukey-Kramer test revealed the pressure delivered to the dACC was significantly lower than both AI and PI (p < 0.05) and the pressure delivered to PI was significantly higher than AI **Figure 3**.

**Figure 3.**
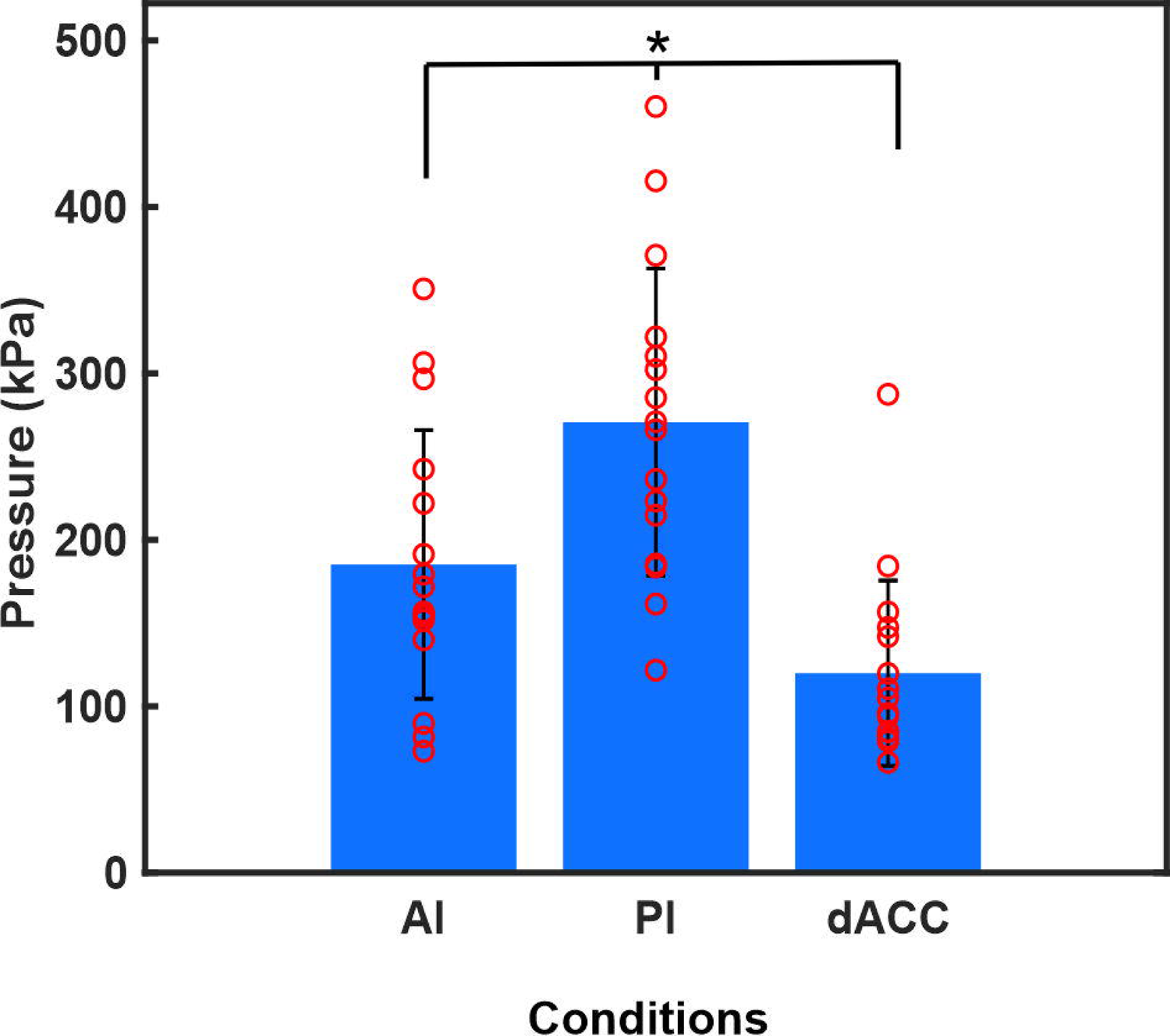
Estimated pressure. Group (N = 16) mean ± SEM estimated pressure in the brain for each of the LIFU conditions. Red circles represent individual subject data. AI = anterior insula; PI = posterior insula; dACC = dorsal anterior cingulate cortex. * = p < 0.05.

### Pain Thresholds

The average pain threshold for AI, PI, dACC, and Sham sessions was: 44.0 ± 4.2 °C, 46.1 ± 3.2°C, 44.5 ± 3.5 °C and 47.0 ± 2.5 °C. A one-way repeated measures ANOVA revealed no differences in pain threshold across sessions: F(3,51) = 0.86, p = 0.47.

The average 5/9 pain rating for AI, PI, dACC, and Sham sessions was: 46.1 ± 3.2 °C, 47.0 ± 2.5°C, 46.3 ± 3.7 °C and 45.8 ± 3.8 °C. A one-way repeated measures ANOVA revealed no significant differences in 5/9 pain rating across sessions: F(3,51) = 1.67, p = 0.19. Individual participant data is provided in **Supplementary Table 2.**

### CPM

#### Conditioning stimulus submerge times

The mean ± SD submerge differences scores (Post-Pre) averaged across the three trials for AI, PI, dACC, and Sham was: −6 ± 35 seconds, 13 ± 19 seconds, 5 ± 19 seconds, and −2 ± 16 seconds. The one-way repeated measures ANOVA was not significant: F(3,45) = 2.31, p = 0.089.

#### Pain Ratings

CPM was conducted 3 times before and after the intervention. Participants were required to rate pain intensity after taking their hand out of the cold water (conditioning stimulus). To account for potential day-to-day differences in pain ratings post-intervention data was subtracted from pre-intervention data. A two-way repeated measures ANOVA was performed with main factors LIFU (AI, PI, dACC, Sham) and TIME (0, 10, 20 seconds) and the LIFU*TIME interaction. The two-way repeated measures ANOVA revealed a main effect of LIFU: F(3,180) = 3.56, p = 0.0155. There was no main effect of TIME: F(2,180) = 0.3, p = 0.74 and there was no interaction of LIFU*TIME: F(6,180) = 0.3, p = 0.94 (**Figure 4A**). Tukey-Kramer post-hoc testing of the main effect of LIFU revealed LIFU to PI was significantly different from AI (p < 0.05) and dACC (p < 0.05) but not Sham (see **Figure 4B**).

**Figure 4.**
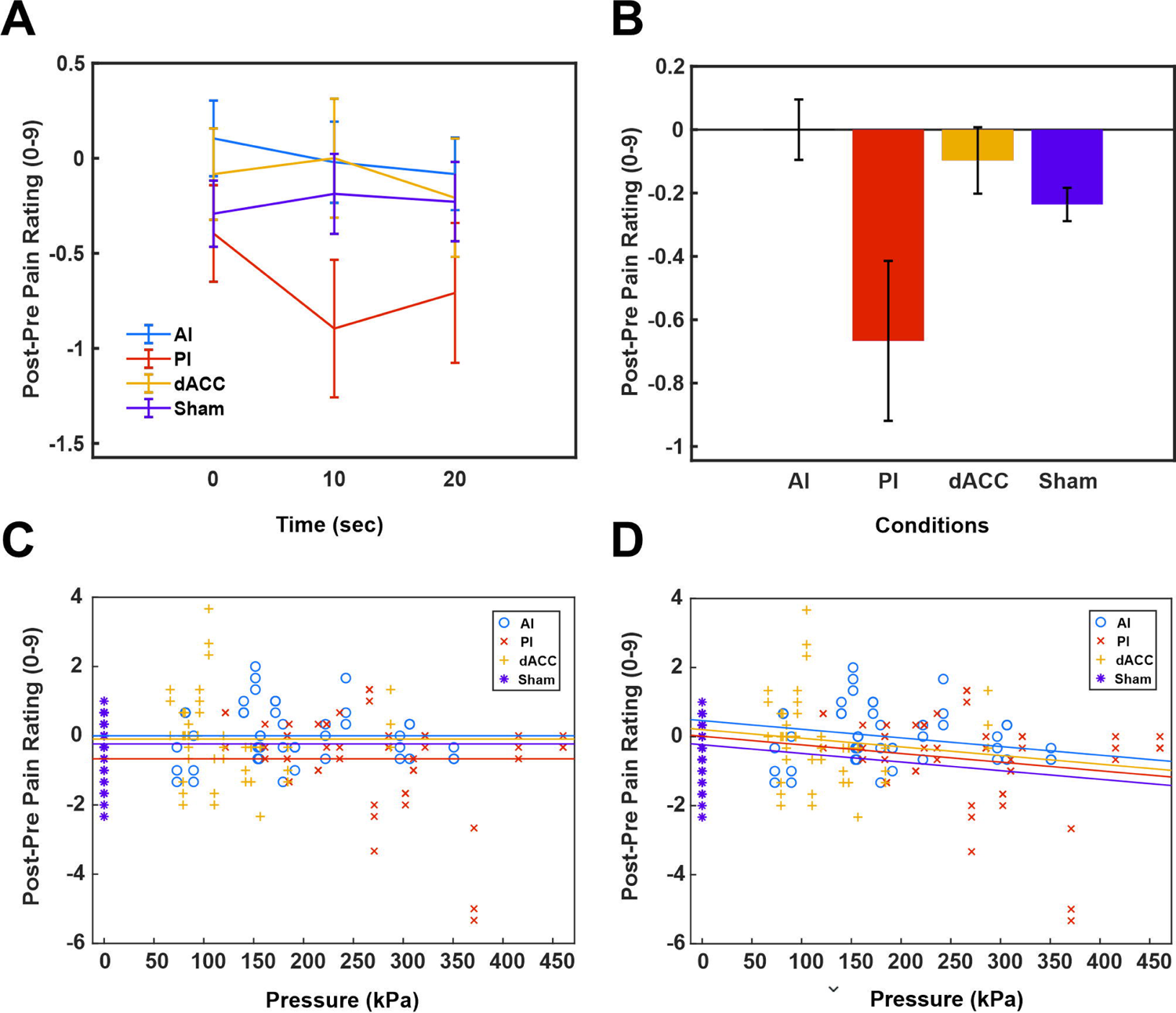
Effects of LIFU on conditioned pain modulation. **A.** Group (N = 16) mean ± SEM pain rating change scores after LIFU (post) subtracted from before LIFU (pre) using a 0-9 point scale. AI = LIFU delivered to anterior insula; PI = posterior insula; dACC = dorsal anterior cingulate cortex; Sham = no LIFU delivered. Time represents timing of the testing stimulus after the conditioning stimulus. 0 = Immediately after. **B.** Group (N = 16) mean ± SEM pain change scores collapsed across time for each of the LIFU targets. **C.** Data from each subject from each time point plotted as a function of pressure in kiloPascals (kPa). Horizontal lines show separate means (same as in B) for each of the conditions. **D.** Data from C with adjusted means showing the effect of accounting for pressure differences between conditions.

Because there was a significant difference in the pressure delivered to each of the brain targets we ran a separate model including pressure as a continuous covariate. To simplify the model we collapsed the data over the time factor to determine the effect of pressure on the group differences (see **Figure 4 C&D**). This analysis revealed a main effect of group F(3,188) = 3.67, p = 0.0134; and when accounting for pressure the group differences were slightly diminished but still significant F(3,185) = 3.16, p = 0.026 (see **Figure 4D**). In addition, we examined if there was a LIFU*PRESSURE interaction to answer the question if there were target specific effects of pressure on pain difference scores. This analysis revealed no interaction F(2,185) = 1.09, p = 0.34. The coefficient estimates for each of the LIFU conditions and the analysis of covariance results of this model is provided in **Supplementary Figure 1**.

### TSP

#### Pain Ratings

The two-way repeated measures ANOVA with main factors LIFU (AI, PI, dACC, Sham) and TIME (0 – 10 seconds) and the interaction LIFU*TIME revealed a main effect of LIFU: F(3,599) = 9.66, p = 0.0001. It also revealed a main effect of TIME: F(9,599) = 2.68, p = 0.0046. The interaction was not significant: F(27,599) = 0.4, p = 0.99. Post hoc tests of the main effect of LIFU revealed PI was significantly lower than both AI (p < 0.05) and Sham (p < 0.05) and that dACC was also significantly lower than Sham (p < 0.05). The post-hoc tests of the main effect of TIME revealed time points 8, 9 and 10 were significantly lower than both time points 1, 2 and 4. Time point 3 was significantly higher than time point 10. All comparisons are p < 0.05. **Figures 5A & B.** To assess the effect of pressure delivered to each of the brain targets we ran a separate model including pressure as a continuous covariate. To simplify the model we collapsed the data over TIME to determine the effect of pressure on the group differences. This analysis revealed a main effect of group F(3,636) = 9.66, p < 0.0001 (**Figure 5C**). When accounting for pressure the group differences were diminished but still significant F(3,635) = 2.81, p = 0.039. (see **Figure 5D**). In addition, we examined if there was a GROUP*PRESSURE interaction to answer the question if there were target-specific effects of pressure on pain difference scores. This analysis revealed no significant interaction F(2,633) = 2.94, p = 0.054. The coefficient estimates for each of the LIFU conditions and the analysis of covariance results of this model is provided in **Supplementary Figure 2**.

**Figure 5.**
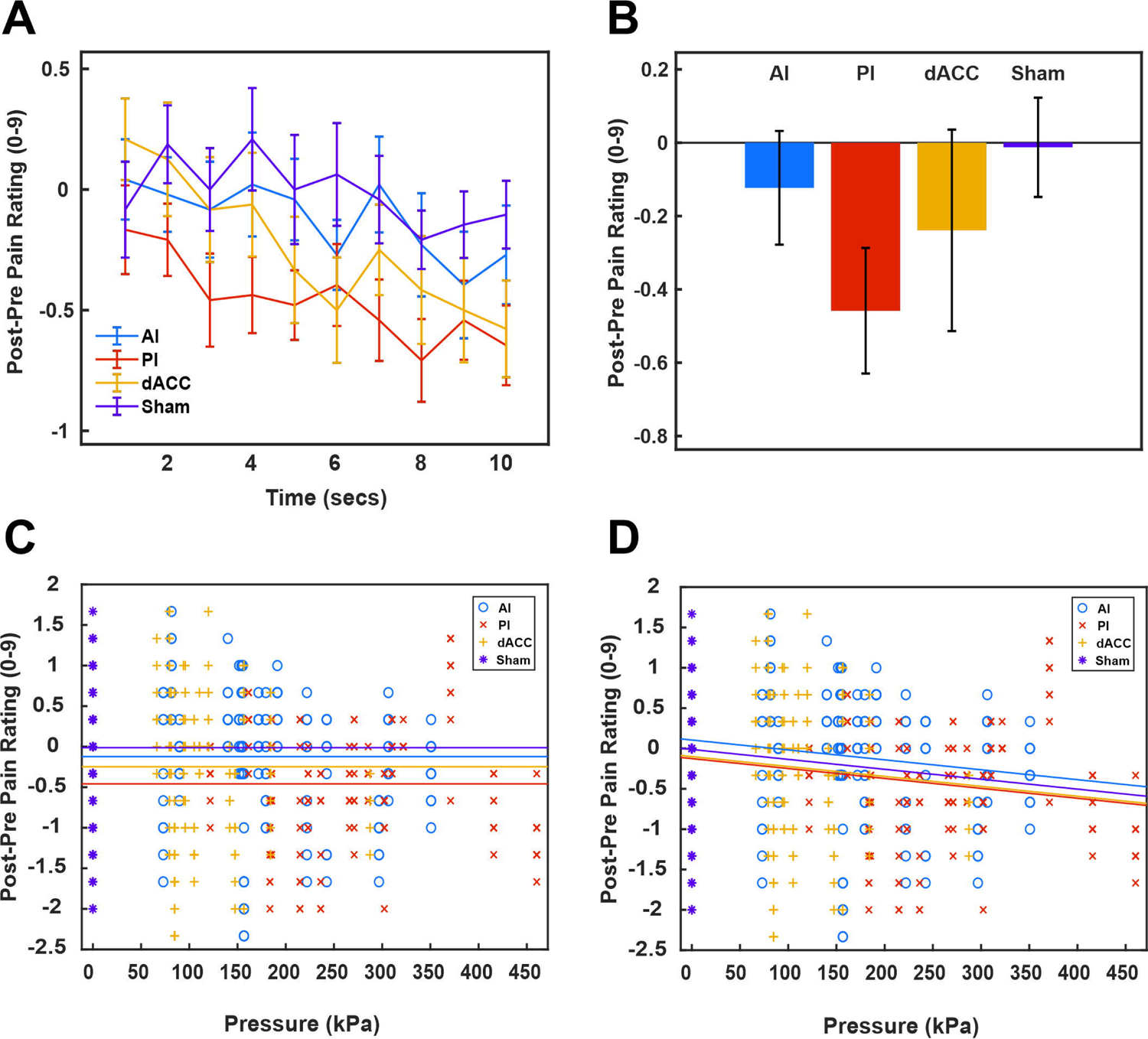
Effects of LIFU on temporal summation of pain. **A.** Group (N = 16) mean ± SEM pain rating change scores after LIFU (post) subtracted from before LIFU (pre) using a 0-9 point scale. AI = LIFU delivered to anterior insula; PI = posterior insula; dACC = dorsal anterior cingulate cortex; Sham = no LIFU delivered. Time represents timing of the testing stimulus after the conditioning stimulus in seconds (sec). **B.** Group (N = 16) mean ± SEM pain change scores collapsed across time for each of the LIFU targets. **C.** Data from each subject from each time point plotted as a function of pressure in kiloPascals (kPa). Horizontal lines show separate means (same as in B) for each of the conditions. **D.** Data from C with adjusted means showing the effect of accounting for pressure differences between conditions.

#### Report of Symptoms

No serious adverse events were reported for any of the interventions. The most common reported symptom both before and after any intervention was sleepiness. The next most common reported symptom before intervention was anxiousness. Instances of anxiousness were reported less frequently after the interventions. Group and individual participant report of symptoms is presented **Figure 6A**.

**Figure 6.**
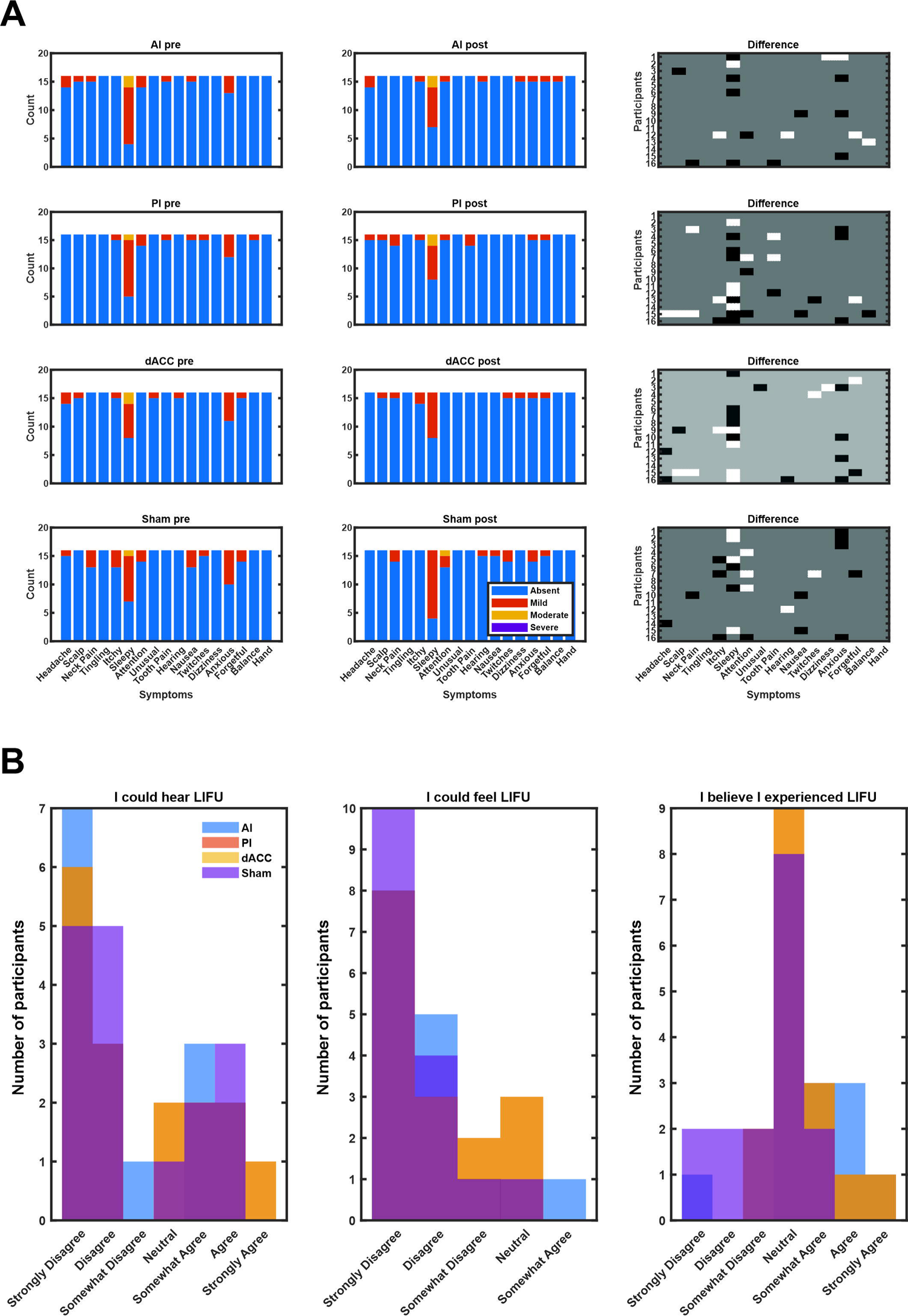
Adverse events and acoustic masking. **A.** Group (N = 16) adverse events data for each condition (rows) before LIFU (pre/first column) and after LIFU (post/second column). The third column represents the difference score (post-pre) for each subject for each queried symptom. White represents an increase while black denotes a decrease in the symptom queried after the intervention. Note in the dACC condition one subject reported a −2 point decrease (black) and all −1 point decreases are dark grey. **B.** Histograms (N = 16) showing the response frequency for the three queried questions color-coded by intervention.

#### Acoustic Masking

The mean ± SD response for the question ‘I could hear the LIFU stimulation’ for AI, PI, dACC, and Sham was: 1.7 ± 2.0, 2.1 ± 2.2, 2.1 ± 2.2, and 1.9 ± 2.0. The median for each was 1 and the mode for each was 0. A Kruskal-Wallis test revealed no significant difference: H(3) = 0.34, p = 0.95.

The mean ± SD response for the question ‘I could feel the LIFU stimulation’ for AI, PI, dACC and Sham was: 0.9 ± 1.2, 1.0 ± 1.2, 1.0 ± 1.2, 0.6 ± 0.9. The median was 0.5, 0.5, 0.5 and 0 respectively and the mode was 0 for all conditions. A Kruskal-Wallis test revealed no significant difference: H(3) = 1.3, p = 0.73.

The mean ± SD response for the question ‘I believe I experience LIFU stimulation’ for AI, PI, dACC and Sham was: 3.6 ± 1.4, 3.4 ± 1.0, 3.4 ± 1.0 and 2.4 ± 1.3. The median and mode for all conditions was 3. A Kruskal-Wallis test revealed a significant effect: H(3) = 8.08, p = 0.044. A post-hoc Dunn’s test revealed this was driven by a significant difference between Sham and AI only. Histograms of the results of the acoustic masking questionnaire are provided in **Figure 6B**. Individual data is shown in **Supplemental Table 3.**

## DISCUSSION

We delivered single-element 500 kHz LIFU to the AI, PI, or dACC as compared to sham for effect on measures of central sensitization including CPM and TSP and examined how pressure affected these measures. LIFU to PI significantly reduced pain ratings as compared to AI, dACC, and Sham during the TSP task and significantly reduced pain ratings during the CPM task compared to AI and dACC only. LIFU to the dACC significantly reduced pain ratings compared to Sham during TSP with no effect during CPM. Despite delivering the same *ex vivo* pressure to each participant to each target site, the estimated intracranial pressure in the brain was significantly different between conditions with the largest pressure delivered to the posterior insula location. As such, we included pressure as a continuous covariate in subsequent statistical models. Including pressure reduced between group differences for both the CPM and TSP task but did not change the overall main effect of LIFU between groups for CPM or TSP.

### Effects of LIFU on central sensitization

Temporal summation of pain refers to the effect of tonic input to C-fibers at a frequency ≥ 0.33 Hz that can induce or result in a central hyperalgesic state sometimes termed windup pain [43]. We delivered tonic contact heat at 0.5 Hz but, on average, did not see evidence for summation of pain over the duration of the 10 stimuli delivered. Variability in response to TSP protocols such that summation is not achieved is well documented [37,39]. Cheng et al. (2015) found that the response to a TSP protocol was correlated with both structural and functional connectivity of primary somatosensory cortex and thalamus as well as cortical thickness in insula. Further, TSP response also looked to be influenced by whether the subjects were considered pronociceptive or antinociceptive [39]. This concept was introduced by Yarnitzky et al. (2014) to profile individuals’ pain processing and morbidity [87]. Pronociceptive individuals have facilitated pain processing and/or reduced descending pain-modulating capacity whereas antinociceptive individuals display the reverse phenotype [39,87]. For example, several chronic pain disorders characterized as having increased central sensitization such as fibromyalgia, irritable bowel syndrome, migraine etc. would have a pronociceptive profile. This study was conducted in pain-free young, healthy individuals and thus this cohort likely fell within an antinociceptive profile, which may explain why robust temporal summation of pain was not achieved. On the group level, there was no significant change in reported pain across stimuli during the sham stimulation but LIFU to PI and dACC significantly reduced pain rating. Both the PI and dACC receive input from lamina I dorsal horn neurons via the spinothalamic tract but through different nuclei in the thalamus, suggesting similar but complimentary roles in bottom-up pain processing [56,58,88]. The PI is the primary cortical terminus for these ascending nociceptive afferents and inhibition of the PI had the largest effect suggesting that our TSP pain rating reduction is likely from bottom-up intensity coding. LIFU to AI did not result in a significant difference from Sham supporting the specific role of PI in bottom-up coding of stimulus attributes such as intensity and for the AI to be more involved in top-down integration of salience and emotional aspects of the pain experience [60,62].

In addition to TSP we also tested conditioned pain modulation as a means of investigating top-down control of pain modulation. Human laboratory tests of CPM are modelled on the concept of ‘pain inhibits pain’ [41] based on mechanisms originally investigated in rats that demonstrated widespread descending inhibition of heterotopic noxious inputs [89]. We originally hypothesized that inhibition of AI would selectively affect CPM as compared to PI due to its purported role in top-down integrative aspects of pain as opposed to bottom-up sensory-discriminative components [60,90,91]. However, LIFU to PI was the only site that demonstrated significant attenuation of pain ratings of the test stimulus. While PI looked to receive the highest estimated intracranial pressure, the intercept and slope for the PI condition was not significantly different from zero suggesting that pressure alone did not account for this effect. Another potential explanation for PI effect is the requirement of the participants was to rate the intensity of the stimulus, of which the PI may subserve a fundamental role independent of any conditioning stimulus employed [92]. Interestingly, the significant effect of PI was only found as compared to AI or dACC stimulation and not Sham stimulation. Under non-pathological conditions, CPM is a normal phenomenon such that pain inhibits pain and is indicative of a properly functioning descending pain control system. This was apparent in our Sham condition where there was a modest decrease in reported pain in this cohort of healthy participants which likely explains why the increase in pain inhibition with PI was not found compared to Sham.

### LIFU dosing and effect of pressure on neuronal activity

There is currently no definitive data on LIFU dosing for neuromodulatory efficacy. Upper limits of dosing or pressure or intensity for neuromodulation are loosely based on the Food & Drug Administration and European International Electro-technical Commission [93] diagnostic safety limits that sets peak average and temporal average limits to avoid potential cavitation and thermal effects. While most LIFU studies adhere to these standards, their relation to neuromodulation effect is poorly understood. Previous human studies have reported estimated intracranial pressure and/or intensity yet relation to effect was not reported [12,94]. Several *in vitro* studies have reported differential effects of pressure [95–98] and in small animals, intensity has been found to relate to motor responses in mice [99–101], including deafened mice models [102]. However, in rabbit, using a blood-oxygen-level dependent (BOLD), doubling intensity did not increase the BOLD response [103] and increasing pressure from 200 kPa to 800 kPa directed at sub-cortical structures in primate did not appreciably affect behavior [104]. While our results are not definitive for the role of pressure towards neuromodulation effect in humans the results demonstrate a contribution of pressure. It is likely that pressure or intensity does not operate in isolation to induce neuromodulation as several other parameters have been theoretically hypothesized [105] and demonstrated to contribute to neuromodulatory response in small animal [4,5,106] and human [13].

## LIMITATIONS

This work has several limitations. For the CPM task, we did not set a controlled time for hand submersion. While conceptually this controls that each person reached a 5/9 pain rating, there was high variability in hand submerge times that could influence levels or whether CPM was achieved. Neither the CPM nor the TSP tasks resulted in the proposed canonical response – reduced pain for CPM and increase pain (over time) for TSP, and hence the results of this study may address the effect of LIFU on pain intensity and not *de facto* central sensitization mechanisms.

## CONCLUSIONS & FUTURE WORK

500 kHz single-element LIFU is an effective non-invasive method to target sub-regions of the insula and the dACC that demonstrates site-specific effects on behavioral measures of pain processing. Future work will aim to evaluate the effects of LIFU on central sensitization in chronic pain populations as well as determine methods to protract pain reduction.

---

**Supplemental Figure 1.**
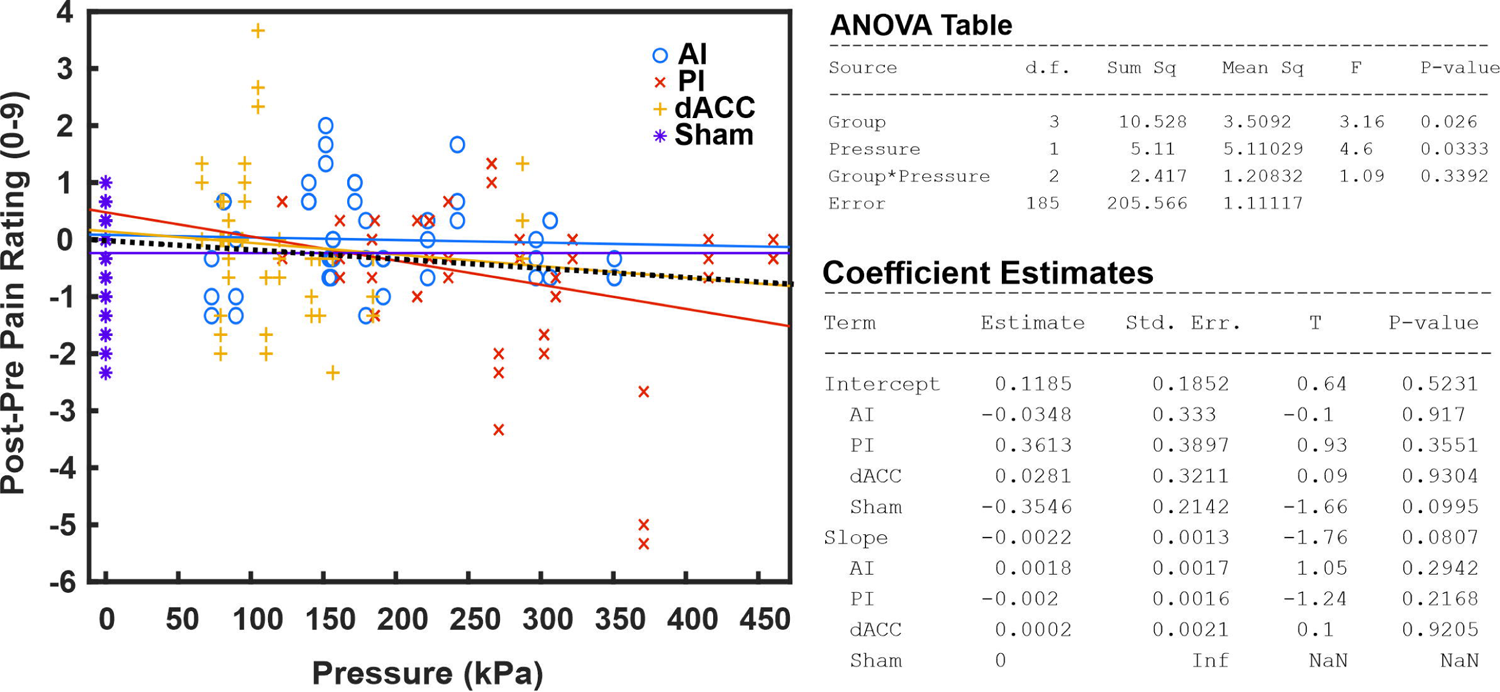
Effects of pressure on each intervention target for condition pain modulation. Figure shows pain change scores as a function of estimated pressure with separate modelled regression lines for each condition. ANOVA table and coefficient estimates from this model are shown on the right.

**Supplemental Figure 2.**
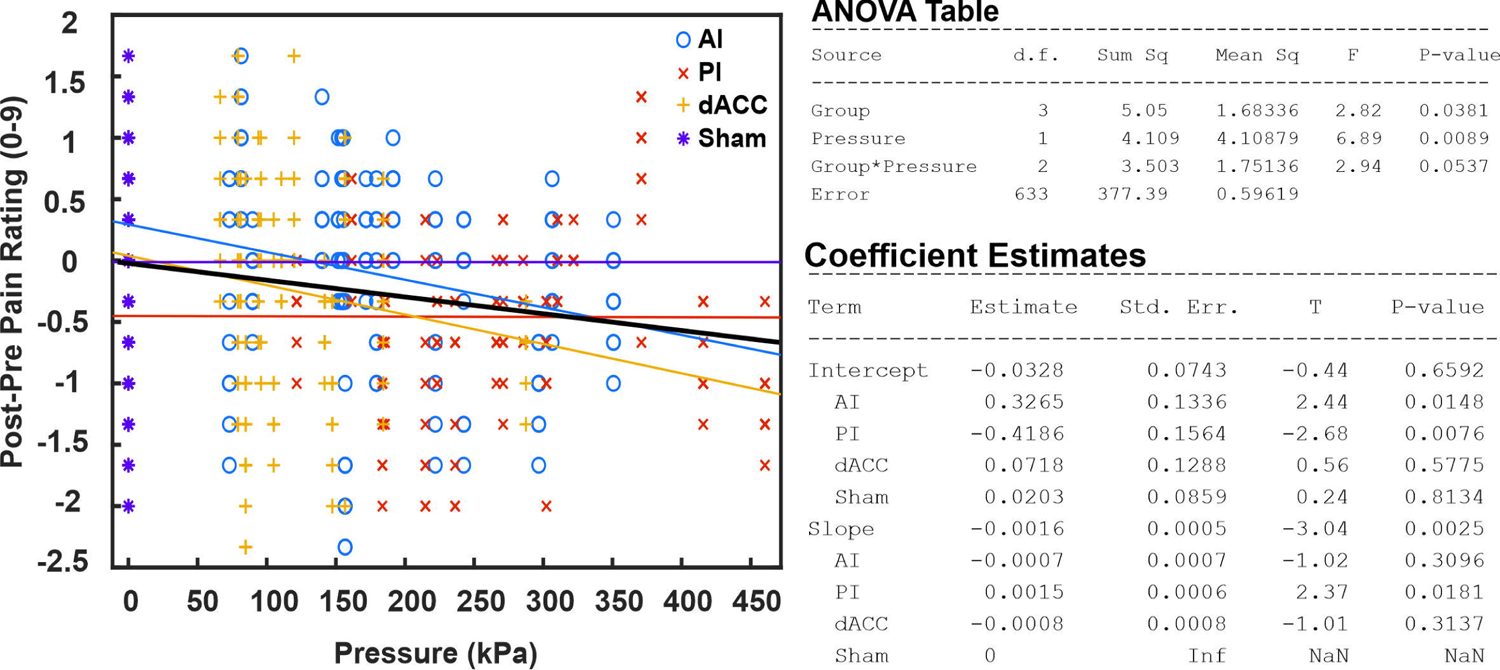
Effects of pressure on each intervention target for temporal summation of pain. Figure shows pain change scores as a function of estimated pressure with separate modelled regression lines for each condition. ANOVA table and coefficient estimates from this model are shown on the right.

