## Supplemental Table 1 for "Low-intensity focused ultrasound to the insula and dorsal anterior cingulate has site-specific and pressure dependent effects on pain during measures of central sensitization"

|  |  | **MNI Coordinates** | | | | | | **Target Depth (mm)** | | |
| --- | --- | --- | --- | --- | --- | --- | --- | --- | --- | --- |
|  |  | **AI** | | | **PI** | | | **AI** | **PI** | **dACC** |
|  |  | **X** | **Y** | **Z** | **X** | **Y** | **Z** |  |  |  |
| **Male** | S1 | -31 | 22 | -3 | -39 | -6 | 4 | 40.8 | 42.4 | 49.3 |
|  | S2 | -33 | 21 | -6 | -36 | -10 | -1 | 37.7 | 45 | 53.7 |
|  | S3 | -38 | 21 | 1 | -39 | -3 | 2 | 30.4 | 35.8 | 55.3 |
|  | S4 | -33 | 14 | 0 | -34 | -12 | 4 | 35.8 | 42 | 45.6 |
|  | S5 | -33 | 24 | -3 | -36 | -11 | 2 | 35.8 | 40.7 | 47.8 |
|  | **Mean** |  | | | | | | **36.1** | **41.2** | **50.3** |
|  | **SD** |  | | | | | | **3.8** | **3.4** | **4.1** |
| **Female** | S6 | -31 | 16 | -1 | -34 | -14 | 1 | 36.3 | 41.9 | 48.8 |
|  | S7 | -33 | 23 | -1 | -35 | -4 | -1 | 34.5 | 38.9 | 46 |
|  | S8 | -37 | 21 | -6 | -39 | -6 | -4 | 33.6 | 36.9 | 47 |
|  | S | -33 | 21 | -3 | -34 | -4 | 1 | 31.2 | 37.3 | 43.8 |
|  | S10 | -33 | 19 | -4 | -36 | -10 | -1 | 33.4 | 38.1 | 46.7 |
|  | S11 | -35 | 20 | -2 | -37 | -9 | -1 | 34.7 | 39.7 | 45.7 |
|  | S12 | -34 | 18 | -4 | -35 | -5 | 2 | 34.9 | 40.3 | 45.9 |
|  | S13 | -34 | 19 | -5 | -33 | -10 | 3 | 36.2 | 41.2 | 45.7 |
|  | S14 | -34 | 21 | -4 | -35 | -7 | 0 | 34.3 | 38.1 | 42.7 |
|  | S15 | -31 | 20 | -4 | -37 | -8 | 4 | 34 | 37.2 | 41.5 |
|  | S16 | -35 | 17 | -2 | -35 | -7 | 1 | 32.3 | 38.8 | 46.6 |
|  | **Mean** |  | | | | | | **34.1** | **38.9** | **45.5** |
|  | **SD** |  | | | | | | **1.5** | **1.7** | **2.1** |

**Table S2. Insula and dACC targets**
