## Supplemental Table 2 for "Low-intensity focused ultrasound to the insula and dorsal anterior cingulate has site-specific and pressure dependent effects on pain during measures of central sensitization"

|  | **AI** | | **PI** | | **ACC** | | **Sham** | |
| --- | --- | --- | --- | --- | --- | --- | --- | --- |
|  | Onset | 5/9 Pain | Onset | 5/9 Pain | Onset | 5/9 Pain | Onset | 5/9 Pain |
| S1 | 46.8 | 47.3 | 46.7 | 47.9 | 44.1 | 45.3 | 47.3 | 48.7 |
| S2 | 44.5 | 46.2 | 45.8 | 47.3 | 40.5 | 46.2 | 46.3 | 44.5 |
| S3 | 34.6 | 39.9 | 35.1 | 42.3 | 33.1 | 40.9 | 35.1 | 35.2 |
| S4 | 44.2 | 47 | 44.9 | 46.2 | 43.2 | 45.8 | 44.8 | 46.3 |
| S5 | 47.5 | 49 | 46.7 | 47.9 | 45.6 | 49.9 | 47.2 | 48 |
| S6 | 47.9 | 48.1 | 47.9 | 49.2 | 47.3 | 48.3 | 48 | 49.1 |
| S7 | 45.5 | 48.9 | 43.1 | 44.9 | 46.7 | 48.5 | 42.4 | 44.2 |
| S8 | 47.7 | 48.7 | 47.3 | 47.7 | 45.8 | 47.1 | 46.4 | 47.8 |
| S9 | 48.5 | 46.5 | 45.5 | 49.4 | 47.4 | 48.9 | 48 | 47.5 |
| S10 | 35.3 | 38 | 38 | 42.6 | 40.1 | 35.6 | 38 | 39.5 |
| S11 | 43.5 | 47.9 | 46.2 | 48.6 | 46.6 | 48.2 | 46.3 | 47.8 |
| S12 | 42.3 | 44.4 | 45.2 | 47.2 | 42.4 | 46.9 | 45.3 | 48 |
| S13 | 40.8 | 45.4 | 43.5 | 44 | 46.6 | 47.1 | 43 | 44.1 |
| S14 | 47.6 | 49.3 | 47.6 | 51.3 | 47.2 | 50.1 | 46.9 | 50.2 |
| S15 | 44.4 | 44.1 | 44.8 | 47.5 | 42.9 | 44 | 46.2 | 46.3 |
| S16 | 43.2 | 46.8 | 43.6 | 47.4 | 41.8 | 48.5 | 41.2 | 44.8 |
| **Mean** | **44.0** | **46.1** | **44.5** | **47.0** | **43.8** | **46.3** | **44.5** | **45.8** |
| **SD** | **4.2** | **3.2** | **3.5** | **2.5** | **3.8** | **3.7** | **3.7** | **3.8** |
