## Supplemental Table 3 for "Low-intensity focused ultrasound to the insula and dorsal anterior cingulate has site-specific and pressure dependent effects on pain during measures of central sensitization"

|  | **AI** | | | **PI** | | | **ACC** | | | **Sham** | | |
| --- | --- | --- | --- | --- | --- | --- | --- | --- | --- | --- | --- | --- |
|  | **Hear** | **Feel** | **Believe** | **Hear** | **Feel** | **Believe** | **Hear** | **Feel** | **Believe** | **Hear** | **Feel** | **Believe** |
| S1 | 0 | 0 | 3 | 3 | 0 | 3 | 0 | 0 | 3 | 0 | 0 | 0 |
| S2 | 5 | 4 | 5 | 5 | 2 | 4 | 5 | 2 | 2 | 5 | 2 | 2 |
| S3 | 4 | 1 | 3 | 5 | 1 | 3 | 5 | 1 | 3 | 5 | 1 | 3 |
| S4 | 0 | 0 | 5 | 0 | 0 | 6 | 6 | 4 | 6 | 0 | 0 | 3 |
| S5 | 1 | 0 | 3 | 1 | 1 | 3 | 1 | 1 | 3 | 1 | 0 | 3 |
| S6 | 0 | 1 | 3 | 0 | 2 | 3 | 0 | 1 | 3 | 1 | 1 | 1 |
| S7 | 5 | 0 | 4 | 0 | 0 | 3 | 3 | 0 | 3 | 0 | 0 | 3 |
| S8 | 0 | 0 | 6 | 4 | 0 | 5 | 5 | 0 | 6 | 0 | 0 | 3 |
| S9 | 1 | 2 | 3 | 0 | 0 | 4 | 4 | 0 | 3 | 1 | 0 | 3 |
| S10 | 0 | 1 | 4 | 3 | 1 | 2 | 0 | 0 | 4 | 4 | 1 | 4 |
| S11 | 4 | 0 | 3 | 0 | 0 | 3 | 4 | 1 | 4 | 1 | 0 | 4 |
| S12 | 4 | 3 | 4 | 1 | 3 | 2 | 1 | 2 | 2 | 1 | 1 | 1 |
| S13 | 1 | 1 | 3 | 0 | 0 | 3 | 5 | 0 | 3 | 0 | 0 | 3 |
| S14 | 0 | 1 | 5 | 1 | 3 | 4 | 0 | 0 | 3 | 3 | 3 | 2 |
| S15 | 2 | 0 | 0 | 4 | 3 | 3 | 4 | 1 | 0 | 4 | 0 | 0 |
| S16 | 0 | 0 | 3 | 6 | 0 | 3 | 5 | 0 | 3 | 5 | 0 | 3 |
| **Mean** | **1.7** | **0.9** | **3.6** | **2.1** | **1.0** | **3.4** | **3.0** | **0.8** | **3.2** | **1.9** | **0.6** | **2.4** |
| **Median** | **1** | **0.5** | **3** | **1** | **0.5** | **3** | **4** | **0.5** | **3** | **1** | **0** | **3** |
| **Mode** | **0** | **0** | **3** | **0** | **0** | **3** | **5** | **0** | **3** | **0** | **0** | **3** |
| **SD** | **2.0** | **1.2** | **1.4** | **2.2** | **1.2** | **1.0** | **2.3** | **1.1** | **1.4** | **2.0** | **0.9** | **1.3** |
